# An automated morphometric approach to evaluate distal lung patterning in mouse models of bronchopulmonary dysplasia

**DOI:** 10.1101/2025.09.11.675692

**Authors:** Declan J. Gainer, Mark L. Ormiston

## Abstract

**Background:** Chronic respiratory diseases represent a large group of non-communicable diseases that are a leading cause of mortality and morbidity globally. Many of the methods utilized to assess airway simplification in experimental models of the conditions are overly time-consuming and are sensitive to inter-operator biases, necessitating the need for unbiased and efficient tools to supplement analyses.

**Methods:** We propose a semi-automated method to quantitate the characteristics of large terminal respiratory airways and alveoli that uses free image-processing software (Fiji). We aimed to develop and test this method in a mouse model of bronchopulmonary dysplasia (BPD), a disease of blunted airway and pulmonary vascular development that remains a leading cause of mortality among preterm infants. Optimal macro parameters were determined with a test set of images from postnatal day 14 (P14) mice exposed to acute postnatal hyperoxia by determining which area and circularity values best correlated with mean linear intercept (L_M_). Validation was performed on a separate set of images from P7 mice subjected to the same hyperoxic model of BPD.

**Results:** Both alveolar duct (r: 0.7866, p = 0.0359) and alveolar (r: 0.9475, p = 0.0012) area correlated with L_M_ measurements from the test set. Using our method on a validation dataset, we demonstrate that hyperoxia-exposed mice possess fewer, enlarged alveoli that occupy less total area, as well as enlarged alveolar ducts that occupy a greater proportion of the parenchyma.

**Conclusions:** We report a semi-automated method of quantitating the characteristics of large and small terminal respiratory airways. This tool expedites analysis and removes operator bias relative to existing methods. We also demonstrate that L_M_ changes in the acute hyperoxia-induced BPD model result from both alveolar simplification and inadequate primary septation at the level of the alveolar ducts.

## Introduction

Many disorders are known to negatively influence the lung parenchyma and airways. Collectively, these disorders are known as chronic respiratory diseases (CRDs) (1), a group of non-communicable diseases that remain a leading cause of morbidity and mortality globally (2). CRDs encompass both obstructive and restrictive lung diseases, which are known to manifest differing structural phenotypes as a consequence of underlying disease pathophysiology and environmental influences. Classically, obstructive diseases are characterized by the presence of emphysema-like structural changes, or the enlargement of distal airspaces due to the destruction of parenchymal tissue (3). This enlargement can affect entire pulmonary lobules (panlobular), or can be primarily localized to the respiratory bronchioles (centrilobular) (4, 5). In comparison, restrictive diseases are driven by the accumulation and deposition of excess fibrotic tissue throughout the lung, in addition to matrix-related proteins such as elastin and collagen (6). These structural changes not only impede gas exchange across the respiratory membrane, but also thicken airway walls, reducing their compliance and functional diameter (6).

Although lung diseases are generally be classified as either obstructive or restrictive in nature, bronchopulmonary dysplasia (BPD) can exhibit characteristics of both, making it a phenotypically diverse disease (7). BPD is a chronic lung disease of prematurity that has been linked to the blunted development of both the pulmonary circulation and airways, resulting in persistent airway dysfunction and an increased risk of various cardiopulmonary complications later in life (8). This disease has been associated with a range of antenatal and postnatal risk factors, including pre-term birth and early-life respiratory support, and remains a leading cause of mortality and morbidity in preterm infants for which no curative treatments are clinically available. Rather, current therapies focus primarily on minimizing lung damage. Consequentially, experimental approaches to treat BPD are often tested in animal models, commonly mouse models of BPD, which typically involve postnatal exposure to hyperoxia (75%-95% O_2_) for an extended duration, ranging from 3-14 days (9). Models utilizing invasive ventilation, mixed hyperoxia and hypoxia, chorioamnionitis, and transgenic modifications have also be utilized.

Although experimental timelines and exposures are variable, continuous exposure to hyperoxia is known to blunt lung development during the late saccular and early alveolar stages of lung development. Much like in humans, this results in alveolar simplification, fibrosis, and the development of a dysregulated pulmonary vasculature (9). One of the challenges of these models are the methods used to quantify the impact of disease on airway patterning. Several methods are conventionally used to quantify these impacts, including radial alveolar counts (RAC) (10), mean linear intercept (L_M_) (11), and gold-standard lung stereology (12). These methods aim to provide quantitative measures of structural aspects of the distal lung, including airway size (L_M_), alveolar septation (RAC), and absolute alveolar number and surface area (stereology). Although robust and commonplace, all three methods are time-consuming, particularly stereology, due to specific requirements for inflation, embedding, sectioning, and imaging. Additionally, operator bias and a lack of specificity for specific generations of airways are considerations, highlighting the need for time-efficient, unbiased tools that can quantitate the characteristics of different classes of airways. Here, we propose a tool to supplement these conventional measures in assessing disease in a mouse model of BPD that is capable of providing both airway counts and area measurements in a manner that reduces assessment time and circumvents operator bias.

## Materials & Methods

### Ethics Statement

All animal work was performed at Queen’s University in Kingston, Ontario, Canada. Experimental protocols (#2024-2223) were approved by the University Animal Care Committee (UACC) and adhered to guidelines established by the Canadian Council on Animal Care (CCAC). Mice were housed in individually ventilated cages and received *ad libitum* access to food and water.

### Mouse model of bronchopulmonary dysplasia

Timed-pregnant C57BL/6 dams (Charles River Laboratories) were kept at room temperature on 12 h sleep/wake cycles until birth. Following birth on postnatal day 0 (P0), all newborn pups were assigned to (i) normoxia (21% O_2_) or (ii) hyperoxia (95% O_2_) groups. Pups in group (ii) and their respective dams were exposed to hyperoxia in a single-latch A-Chamber (BioSphreix) for 3 days (13), while pups in group (i) were housed in the same room under normoxic conditions. Dams were rotated pairwise between conditions every 24 hours to prevent oxygen toxicity and preserve maternal health. Pups were removed from 95% O_2_ after 72 hours on P3, and were allowed to recover under normoxic conditions until either P7 or P14. A subset of neonates received a subcutaneous dose of 5 µL PBS prior to hyperoxic exposure. Animals were euthanized via an intraperitoneal overdose of Sodium Pentobarbital (100 mg/kg), in accordance with CCAC guidelines.

### Lung tissue harvest and processing

Following sacrifice, the chest cavity was opened, and the right ventricle was perfused with heparinized saline (1000 U/mL diluted 1:50 in phosphate buffered saline (PBS)). The lungs were inflated with 2% paraformaldehyde (PFA) at a pressure of 25 mmHg, were removed *en-bloc*, and were subsequently fixed in 2% PFA at 4°C for 48 hours in the dark. Lungs were subsequently washed overnight in 1X PBS at 4°C, embedded in paraffin, sectioned (5 µm), and stained with hematoxylin and eosin (Abcam). Slides were imaged with a MICA (Leica Microsystems). Six (6) 20X high-powered fields of view (HPF) were taken per animal.

### Manual mean linear intercept scoring

L_M_ scores were obtained from Fiji by overlaying a line of known length over a region of distal lung tissue, avoiding terminal respiratory bronchioles or vascular structures, and counting the number of intersections with alveolar walls. Scores were calculated as:

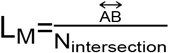

where 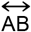 is the length of the overlaid line, and N_intersection_ is the number of times a given line intersected with an alveolar wall. Two measurements were taken per HPF by a single blinded observer, totalling 12 measurements per animal which were averaged to compute the final L_M_ score.

### Method workflow

Briefly, a Fiji (14) macro was written to provide a semi-automated method of quantitating the characteristics of distal airways, specifically alveolar ducts and alveoli. Obtained measurements, including tissue area, airspace area (individual and total), number, and percent of total area, are directly written to an excel file using an existing excel extensions repository (15). Disconnected vascular exudates and small holes (<50 µm^2^) are automatically removed and filled during image processing. Connected exudates, terminal respiratory bronchioles, and major vascular structures must be manually selected and deleted to prevent inclusion.

A user-defined scale must be set globally prior to use, to prevent inaccuracies in output values. Images of most formats and sizes will be accepted, though the macro was developed and tested using.tif image files. Inputs are processed and saved to an output directory for post-processing visual inspection.

The workflow for the method applies to all images being processed sequentially. Following the definition of input and output directories for both test images and the results datasheet, a calibration image is opened from the defined input directory. Here, a user-defined threshold can be set. This threshold assumes all images in the test set are equivalently stained. Opened 24-bit RGB images are then converted to 8-bit greyscale by isolating the green channel (**Fig. 1A,B**). Green was selected to provide superior contrast between stained tissues and the background (16). Users are then prompted to manually select and remove exudates, vascular structures, respiratory bronchioles, pleural spaces, and other unwanted regions from analysis using the wand tool (**Fig. 1C**). Images are then automatically thresholded using user-defined values from calibration (**Fig. 1D**). Disconnected exudates, debris, and particulates are removed to minimize inaccuracies (**Fig. 1E**). The “open” binary operator is applied to all images. Briefly, erosion and dilation are performed in succession to remove isolated pixels and smooth the image (**Fig. 1F**). Small holes (<50 µm^2^) in the lung parenchyma are removed to improve visualization (**Fig. 1G**). This operation does not impact the resulting measurements, as all quantified structures are greater than this size. The characteristics of alveolar ducts and alveoli are quantified (**Fig. 1H,I**), and results are written to an excel file in the specified output directory. All output images are automatically saved for visual inspection.

**Figure 1.**
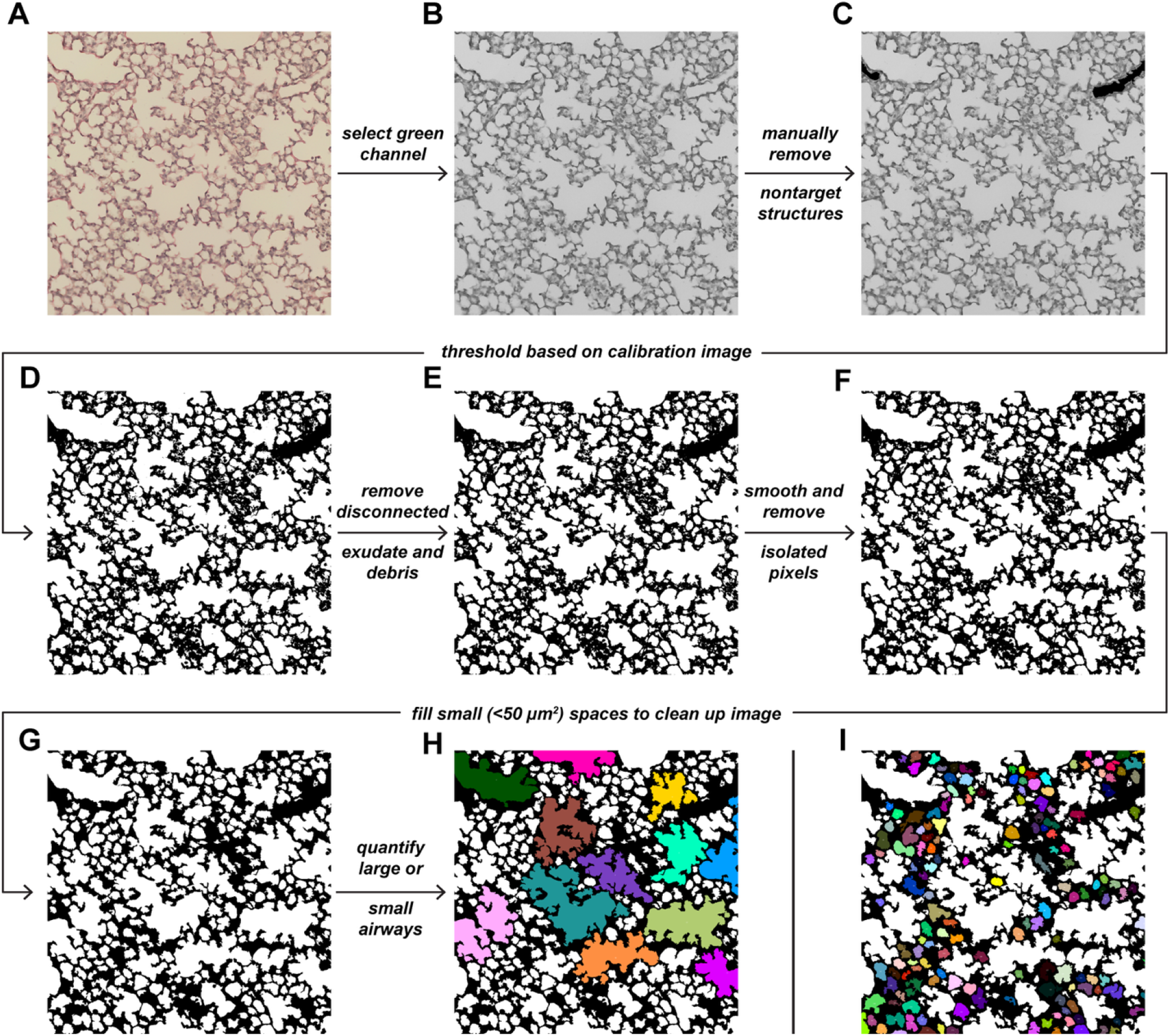
Summary of image processing and analysis steps for quantifying alveolar duct and alveolar area. (**A**) Representative RGB color image acquired at 20x magnification. (**B**) Isolated green channel of **A.** (**C**) Vessels removed from **B**. (**D**) Binarized version of **D**. (**E**) Disconnected exudate and debris removed from **E**. (**F**) Smoothed version of **F**. (**G**) Small holes removed from **G**. Selective quantification of (**H**) alveolar ducts or (**I**) alveoli.

### Method optimization

The irregularity in both shape and size of terminal respiratory airspaces necessitated optimization of the particle analyzer’s parameters. As L_M_ is a measurement that is dependent on airspace size, we sought to identify the parameters that best correlated with manually-derived L_M_ scores by evaluating several combinations of area and circularity values (Tables 1 & 2) using a dataset consisting of lung images from 7 animals. Animals from both healthy and diseased groups were included to ensure a range of L_M_ values were represented. In this context, area refers to target airspace area, and circularity refers to the circularity of target airways.

**Table 1.** Base area and circularity values tested for the quantification of alveolar ducts.

| Airspace Area | 3000 $\mu\text{m}^2$ | | 4000 $\mu\text{m}^2$ | | 5000 $\mu\text{m}^2$ | | 6000 $\mu\text{m}^2$ | |
| --- | --- | --- | --- | --- | --- | --- | --- | --- |
| Circularity | 0-0.5 | 0-1 | 0-0.5 | 0-1 | 0-0.5 | 0-1 | 0-0.5 | 0-1 |

**Table 2.** Base area and circularity values tested for the quantification of alveoli.

| Base Area | 50 µm<sup>2</sup> |  |  | 100 µm<sup>2</sup> |  |  | 150 µm<sup>2</sup> |  |  |
| --- | --- | --- | --- | --- | --- | --- | --- | --- | --- |
| Circularity | 0.01-1 | 0.1-1 | 0.25-1 | 0.01-1 | 0.1-1 | 0.25-1 | 0.01-1 | 0.1-1 | 0.25-1 |

The macro was applied to the same test images that underwent manual L_M_ quantification using the above parameter combinations. To determine the optimal combination for each respiratory airway type, mean duct area and alveolar area measurements from each animal in the optimization dataset were correlated with L_M_ values using an adapted leave-one-out (LOO) approach (17), reporting Pearson’s r as the evaluation metric. A LOO approach was applied to optimization correlation analyses to reduce the influence of any single observation on reported r values, enabling more robust assessment of association in the context of a small *n*. Briefly, each sample from the *n* = 7 optimization dataset was iteratively excluded, and Pearson’s correlation between airway area and L_M_ was computed on the remaining data (*n* = 6). The median r coefficient across all iterations was compared to the r coefficient derived from the correlation obtained using all 7 samples to determine which parameter combination most strongly and consistently correlated with L_M_. As L_M_ and area report values on different unit scales, both measurements were standardized using z-scoring to enable the assessment of proportional, but not fixed, bias and relative agreement by Bland-Altman analysis:

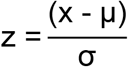

Where x denotes the observed value, µ denotes the variable mean, and σ is the standard deviation of the variable.

### Method validation

Following the determination of optimal circularity and area values, the macro was tested on a separate test set of images from 22 animals. Outputs were compared to manually computed L_M_ scores to determine whether the two measures correlated. Proportional bias and relative agreement were assessed by constructing Bland-Altman plots of standardized measurements, as described above.

### Statistical analysis

All data were collated in GraphPad Prism 10 (v10.6), which was used for statistical analysis. All data are presented as mean ± SD. Assessment of significance between two groups was performed by unpaired, two-tailed Student’s *t*-test. An *f*-test was used to assess differences in variance between experimental groups. If variances differed significantly, Welch’s correction was applied. Pearson’s correlation was used to assess the relationship between L_M_ and alveolar duct/alveolar area. Prior to computing Pearson’s r, normality was assessed for all data using a Shapiro-Wilk test. Linear relationships between L_M_ and duct or alveolar area were confirmed with a scatterplot.

## Results

### Method optimization

To explore the potential for a macro to quantitate the characteristics of distal airspaces, we exposed newborn C57BL/6J mice to 95% O_2_ as an acute model of hyperoxia-induced BPD. This mouse model of BPD arrests lung development as a consequence of hyperoxic exposure, leading to terminal airspace enlargement that can be visualized by histology. In keeping with literature, terminal airspace enlargement was apparent in P14 mice exposed to 95% O_2_ for 72 hours at birth, relative to 21% O_2_ controls, as quantified by L_M_ (**Fig. 2A,B**). To develop a method capable of quantitating the characteristics of distal respiratory airways, and differentiating between alveolar ducts and alveoli, a set of minimum area and circularity values to identify either alveolar ducts or alveoli were selected. A positive association between measured alveolar duct area and L_M_ was observed across all tested parameter combinations (r: 0.7413-0.7922; **Fig. 2C,D**). However, examination of the median and range of r values for the LOO iterations revealed that selecting for the largest alveolar ducts led to less stable associations with L_M_. As such, 4000 µm^2^ - 0-0.5 was selected as the parameter combination that strongly and consistently correlated with L_M_ (r: 0.7866, p = 0.0359; **Fig. 2E**). Bland-Altman analysis of duct area and L_M_ z scores revealed no observations outside the 95% limits of agreement (LOA; 95% LOA: −1.280, 1.280), and no apparent proportional bias (**Fig. 2F**). Similarly, a positive association was observed between measured alveolar area and L_M_ across the tested parameter combinations (r: 0.8516-0.9465; **Fig. 2G,H**). Although the median and range of r values across LOO iterations was consistently higher for alveolar area relative to duct area, 150 µm^2^ - 0.01-1 correlated most stably and strongly with L_M_ (r: 0.9475, p = 0.0012; **Fig. 2I**). Bland-Altman analysis revealed no proportional bias, and no observations fell outside the 95% LOA (−0.6350, 0.6350).

**Figure 2.**
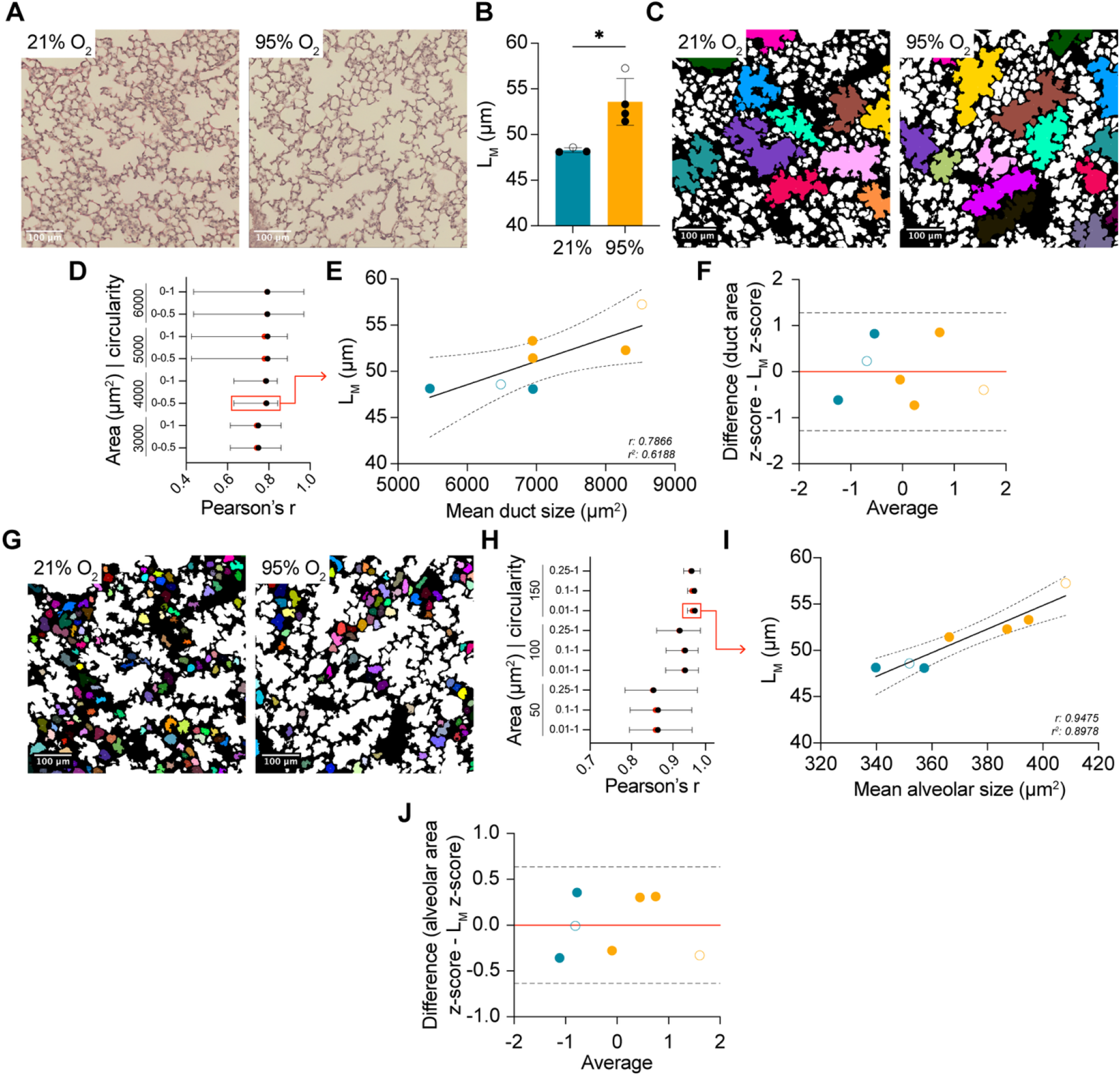
Optimizing method parameters on images from mice exposed to an acute hyperoxia-induced model of BPD. (**A**) Representative H&E-stained lung images from P14 mice exposed to 21% O_2_ (*n* = 3) or 95% O_2_ (*n* = 4) for 72 hours. (**B**) Quantification of mean linear intercept (L_M_) in the mice from **A.** (**C**) Representative colored images depicting alveolar ducts. (**D**) Evaluation of Pearson’s r across minimum area and circularity combinations for alveolar ducts. Red dots depict the r coefficient for all 7 tested samples. Black dots depict the median r coefficient for all 7 LOO iterations. Bars represent the range of values across all 7 LOO iterations. (**E**) Scatter plot of mean duct area plotted against measured L_M_. Solid black line depicts the fitted linear regression line. Dashed lines represent the 95% confidence intervals. (**F**) Bland-Altman plot of z-scores for alveolar duct area and L_M_. Dashed lines represent the 95% limit of agreement (LOA). Red line depicts bias. (**G**) Representative colored images depicting alveoli. (**H**) Evaluation of Pearson’s r across minimum area and circularity combinations for alveoli. Red dots depict the r coefficient for all 7 tested samples. Black dots depict the median r coefficient for all 7 LOO iterations. Bars represent the range of values across all 7 LOO iterations. (**I**) Scatter plot of mean alveolar area plotted against measured L_M_. Solid black line depicts the fitted linear regression line. Dashed lines represent the 95% confidence intervals. (**J**) Bland-Altman plot of z-scores for alveolar area and L_M_. Dashed lines represent the 95% limit of agreement (LOA). Red line depicts bias. (○) female mice, (**•**) male mice. *P < 0.05. Student’s t-test used in **B**. Simple linear regression used in **E, I**. Pearson’s correlation used in **D,H**. Bland-Altman analysis used in **F,J**. Error bars are mean ± SD.

### Method validation

To assess the efficacy of the selected area and circularity parameters in quantitating the characteristics of alveolar ducts and alveoli, the optimized macro was applied to lung images from neonatal mice at P7, following exposure to a model of hyperoxia-induced model of BPD. Although mice from the validation set were at an earlier stage of alveolar lung development than those from the optimization set, acute exposure to 95% O_2_ manifested a similar increase in L_M_ versus controls at P7 (**Fig. 3A,B**). Consistent with this finding, diseased mice had comparatively enlarged alveolar ducts that were more numerous and occupied a greater total area (**Fig. 3D,E,F**). In keeping with previous findings, there was a strong positive relationship between mean alveolar duct area and L_M_ (r: 0.7651, p < 0.0001; **Fig. 3G**) and no apparent proportional bias (**Fig. 3H**). However, one sample from the BPD group fell outside the limits of agreement (95% LOA: −1.344, 1.344). Interestingly, the relative agreement between mean duct area and L_M_ was poorer for the samples from the BPD cohort when compared to normoxic controls, though this is likely attributable to variances in disease phenotype.

**Figure 3.**
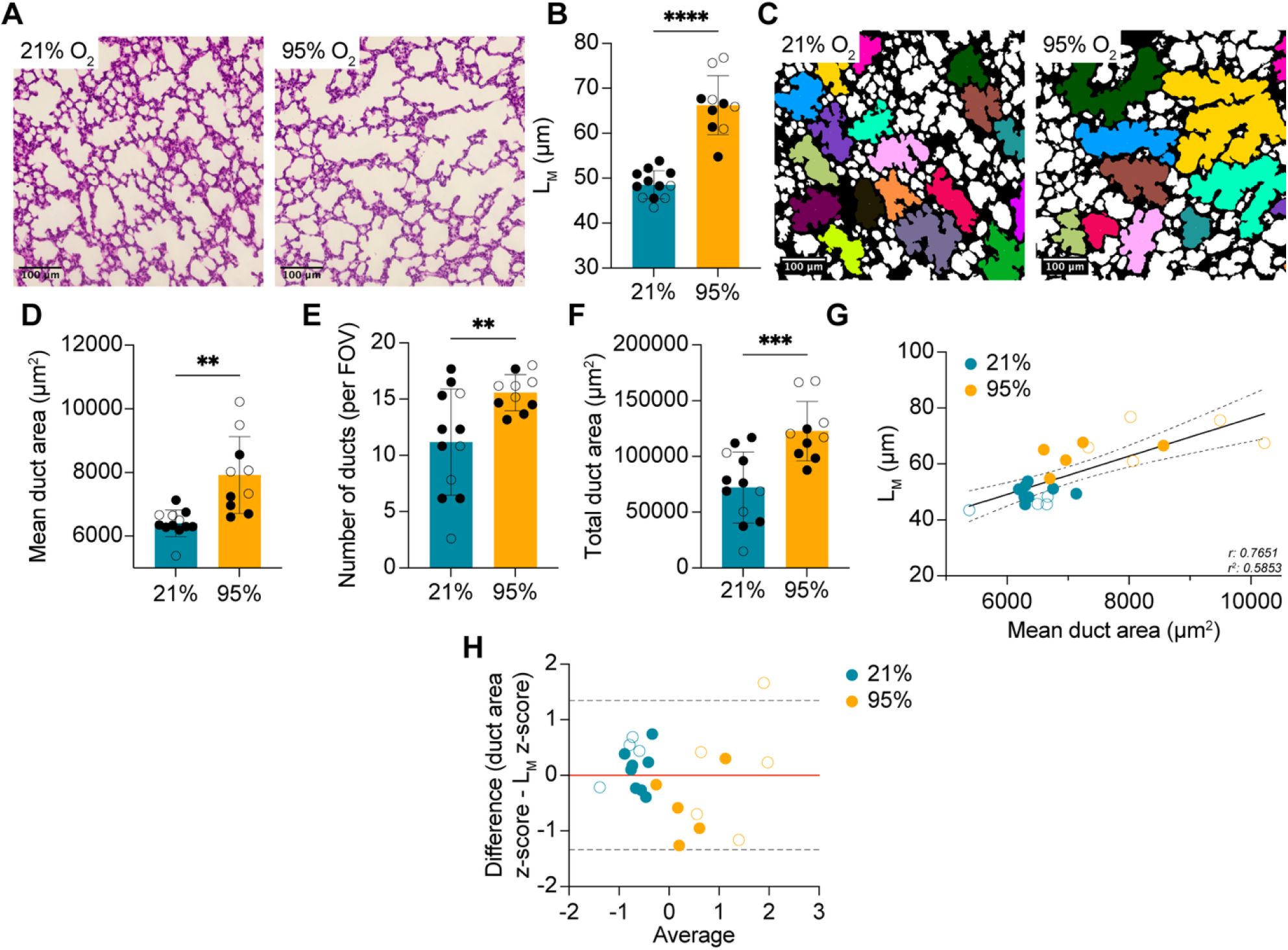
Exposure to postnatal hyperoxia increases alveolar duct size and number. (**A**) Representative H&E-stained images from mice exposed to 21% O_2_ (*n* = 12) or 95% O_2_ (*n* = 10) for 72 hours. (**B**) Quantification of L_M_ in the mice from **A.** (**C**) Representative colored images depicting alveolar ducts. (**D**) Quantification of mean alveolar duct area from the same mice as **A**. (**E**) Quantification of the number of ducts per 20x FOV in the same mice as **A**. (**F**) Quantification of total alveolar duct area in the same mice as **A**. (**G**) Scatter plot of mean alveolar duct area plotted against measured L_M_. Solid black line depicts the fitted linear regression line. Dashed lines represent the 95% confidence intervals. (**H**) Bland-Altman plot of z-scores for alveolar area and L_M_. Dashed lines represent the 95% limit of agreement (LOA). Red line depicts bias. (○) female mice, (**•**) male mice. ****P < 0.0001, ***P<0.001, P** < 0.01. Student’s t-test used in **B,D,E,F**. Simple linear regression used in **G**. Bland-Altman analysis used in **G**. Error bars are mean ± SD.

In keeping with existing literature, our method revealed that the average alveolar size was significantly increased following exposure to acute O_2_ (**Fig. 4A,B**). Additionally, the number of alveoli per FOV and the total area occupied by alveoli were significantly decreased in mice exposed to the hyperoxic model of BPD (**Fig. 4C,D**). Although the association between mean alveolar size and L_M_ remained significant (r: 0.5823, p = 0.0045), the association was weaker than what was computed on optimization data. No proportional bias was evident, and with the exception of two, the majority of samples remained within the 95% LOA (−1.791, 1.791; **Fig. 4F**).

**Figure 4.**
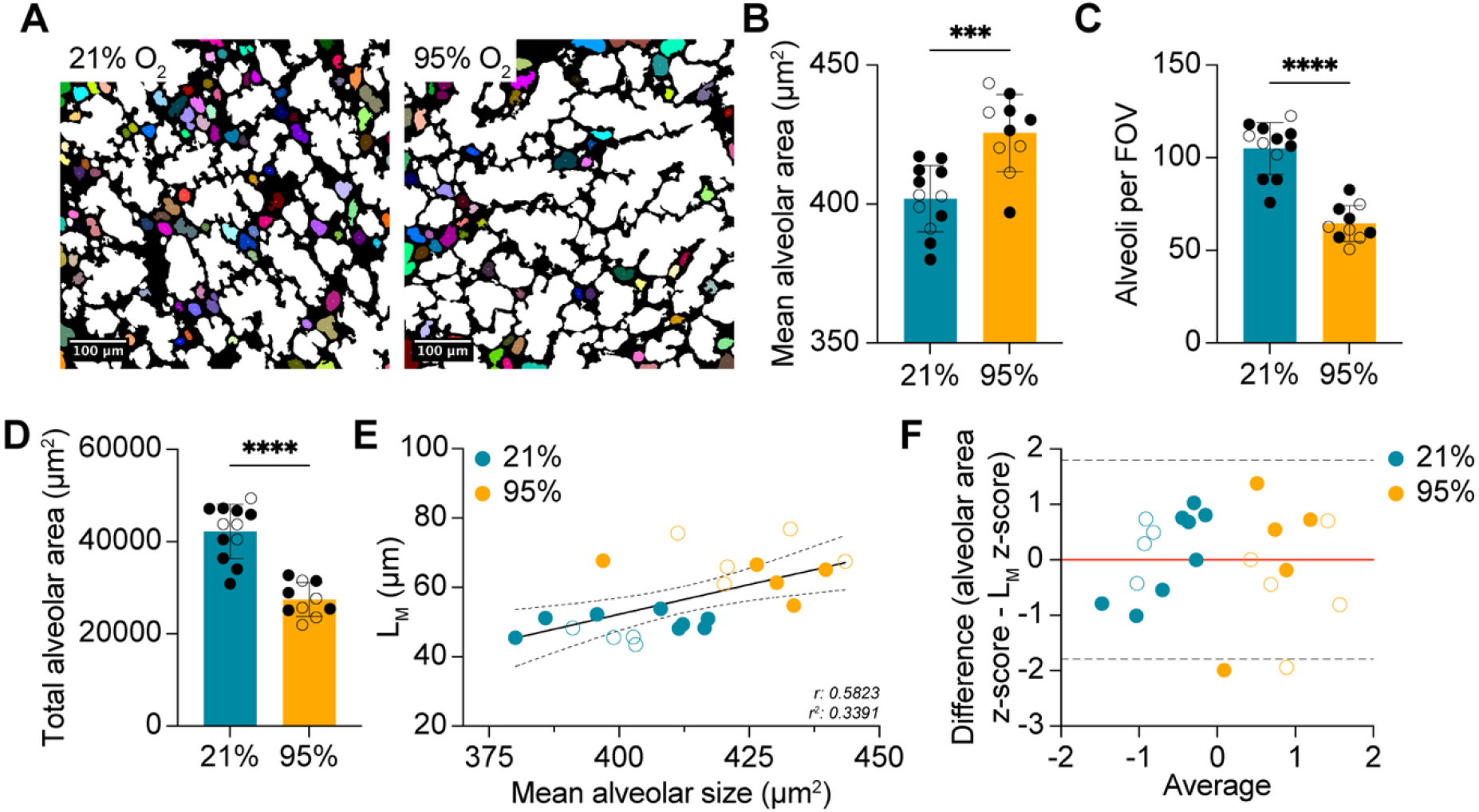
Neonatal hyperoxia impairs alveolar septation. (**A**) Representative colored images depicting alveoli from mice exposed to 21% O_2_ (*n* = 12) or 95% O_2_ (*n* = 10) for 72 hours. (**B**) Quantification of mean alveolar area from the same mice as **A.** (**C**) Quantification of the number of alveoli per 20x FOV in the same mice as **A**. (**D**) Quantification of total alveolar area in the same mice as **A**. (**E**) Scatter plot of mean alveolar duct area plotted against measured L_M_. Solid black line depicts the fitted linear regression line. Dashed lines represent the 95% confidence intervals. (**F**) Bland-Altman plot of z-scores for alveolar area and L_M_. Dashed lines represent the 95% limit of agreement (LOA). Red line depicts bias. (○) female mice, (**•**) male mice. ****P < 0.0001, ***P<0.001, P** < 0.01. Student’s t-test used in **B,C,D**. Simple linear regression used in **E**. Bland-Altman analysis used in **F**. Error bars are mean ± SD.

## Discussion

Here, we demonstrate the utility of a semi-automated method to quantitate the characteristics of both alveolar ducts and alveoli in the distal lung. This method significantly reduces the time required to provide alveolar duct and alveolar counts, while removing the potential for operator bias. Consequently, we propose that this method can be utilized as a means of supporting spatial analyses involving distal lung tissue that use conventional manual measures. Although the optimized set of parameters were validated using tissues from BPD mice versus normoxia-exposed healthy controls, it is likely that this tool can be applied to tissues from other mouse models of lung disease.

While the method is capable of differentiating between alveolar ducts and alveoli, this functionality is dependent on several important assumptions. Firstly, the tool assumes that all large terminal respiratory airways are alveolar ducts, and all small airways are alveoli. This assumption may not be explicitly true, and is dependent on the region of the lung being imaged, orientation, and several other factors that may result in sectioning through a small portion of a large airway. To avoid potential overlap between the two different generations of airways, a buffer zone for minimum area exists to limit the inclusion of intermediate airways that could represent either of the two types. While this buffer may bias the resultant outputs towards slightly smaller alveoli and slightly larger alveolar ducts, the effect is likely minimal, and does not impede the capture of disease effects in our model. Additionally, by allowing for the removal of vascular and nontarget airway structures that may be included in other measurements, we increase the specificity of the results by providing outputs that arise solely from alveolar ducts and alveoli. This feature contrasts with other existing methods of airway assessment that provide related measures, but fail to exclude structures such as large vessels or focus on all airways in the lung, including conducting airways (16, 18).

Inter-observer bias is a consideration for manual methods such as RAC or L_M_. RAC typically involves the placement of a line that sits perpendicular to the center of a respiratory bronchiole. Following placement, the number of alveoli the line transects is counted. This process is repeated several times per lung, and values are averaged per sample. Typically, the main source of inter-observer bias relates to the placement and angle of the perpendicular line. Different observers may choose different angles relative to the pleural space, or different terminal respiratory bronchioles for line placement, a factor that may be influenced by the appearance of said respiratory bronchioles. Although more rigorous, L_M_ measurements are also subjected to related biases, primarily relating to the definition of an intercept. Our method, which aims to supplement these existing measures, provides an unbiased approach to quantitating the characteristics of alveolar ducts and alveoli. In doing so, this tool allows for the determination of whether changes in RAC or L_M_ are driven by changes at the level of one generation of airway, or both, in a manner that eliminates inter-observer bias. Furthermore, the highly automated nature of our method offers significant advantages when supplementing existing morphometric analyses, requiring little additional time or computational power.

While our method minimizes inter-observer bias, it cannot account for bias that results from heterogeneous lung processing (inflation, embedding orientation, sectioning), image acquisition, or the use of improper stereological processing techniques. However, this limitation can be shared by many of the manual morphometric analysis methods, including L_M_ and RAC, and is not the focus of this method.

In conclusion, we have provided an overview of the optimization and validation of a semi-automated macro to characterize alveolar ducts and alveoli in a mouse model of BPD. This method aims to diversify morphometric lung analyses by providing additional measures relating to specific classes of airways, offering insights into whether phenotypic changes are driven by alterations in specific structures. With this method, we demonstrate that mice exposed to acute postnatal hyperoxia develop both enlarged alveoli that are less numerous and occupy a smaller total area, as well as enlarged alveolar ducts. The macro is freely available for use, and is appended as a.ijm macro file.

## Notes

### Competing Interest Statement

The authors have declared no competing interest.

